# Protein pheromone MUP20/Darcin is a vector and target of indirect genetic effects in mice

**DOI:** 10.1101/265769

**Authors:** Sven O. Bachmann, Ellen Cross, Shireene Kalbassi, Matthew Alexandar Sarraf, Michael Anthony Woodley of Menie, Stéphane J. Baudouin

## Abstract

Social behavior in animals is an adaptive process influenced by environmental factors and direct and indirect genetic effects. Indirect genetic effects (IGEs) include mechanisms by which individuals of particular genotypes can influence the behavioral phenotypes and genotypes (via modulated patterns of gene expression) of other individuals with different genotypes. In groups of adult mice, IGEs can be unidirectional, from one genotype to the other, or bidirectional, resulting in a homogenization of the behavioral phenotypes within the group. Critically, it has been theorized that IGEs constitute a large fitness target on which deleterious mutations can have pleiotropic effects, meaning that individuals carrying certain behavior-altering mutations can impose the fitness costs of those mutations on others comprising the broader social genome. Experimental data involving a mouse model support the existence of these IGE-amplified fitness losses; however, the underlying biological mechanisms that facilitate these remain unknown. In a mouse model of IGEs, we demonstrate that the Major Urinary Protein 20 pheromone, also called Darcin, produced by mice lacking the adhesion protein Neuroligin-3 acts as a vector to deleteriously modify the social behavior of wild-type mice. Additionally, we showed that lack of social interest on the part of Neuroligin-3 knockout mice is independent of their environment. These findings reveal a new role for mammalian pheromones in mediating the externalization of social deficits from one individual to others comprising the population through IGEs.

**Author Summary:** Indirect genetic effects (IGEs) are mechanisms by which individuals of particular genotypes can influence the behavioral phenotype of individuals of different genotypes, sometimes disruptively, in instances where one member of the population carriers a deleterious behavior altering variant. Although disruptive IGEs have been demonstrated in mice, its underlying molecular and genetic mechanisms remain unknown. Using an IGEs mouse model, we demonstrated that the pheromone protein Major Urinary Protein 20, also named Darcin, is as a vector and target of social epistasis a specific type of IGEs. This finding reveals a new function for mammalian pheromones in mediating social epistasis to degrade group social behavior.

## Introduction

Social behavior is an adaptive process that environmental factors partly influence [1]. Classical behavior genetic theory posits a dichotomy between genetic and social environmental influences. Experimental work on animal behavior, which has successfully isolated and quantified social environmental and genetic effects, contradicts this theoretical viewpoint and demonstrates that behavior and physiology are dependent on social conditions [2-6]. In fact, the concept of ′social environment′ incorporates indirect genetic effects (IGEs), i.e. those genetic effects arising from the aggregated set of genomes that comprise the organism′s social genotype (or ′social genome′) and which impact the development of an organism′s phenotype and/or genotype [7]. Two types of IGEs have been theorized to exist. *Social genetic effects* capture effects on developing organisms stemming from the action of the social genome on that organism′s phenotype. *Social epistasis,* on the other hand, captures those effects that result from the influence of the social genome on an organism′s genotype (this may take the form of epigenetically modulated patterns of gene expression in phenotypic development) [7,8]. These IGEs are not necessarily unidirectional either, as they could involve the reverse action of individual mutant phenotypes on the social phenotype and genotype also (this will be discussed in more detail below). Molecular evidence for the existence of social epistasis in particular comes from the study of mice, where it has been found that a sizeable portion (as much as 29%) of the variance among mice with respect to certain behavioral phenotypes is the result of altered patterns of gene expression in response to social epistatic transactions with other mice [2]. IGEs might be present in other vertebrate taxa [9-11] and also in invertebrate taxa [6,8,12,13], both in naturalistic and experimental settings [2-6] and may stem from the action of correlational selection of traits during evolution – where the distribution of genes associated with one trait covaries with the genetic variance associated with the distribution of other traits in the population via co-selection – despite the absence of physical patterns of linkage among the genes [14]. A recent model, the *social epistasis amplification model* (SEAM), builds on the concept of social epistasis by further predicting that this phenomenon provides mutations with a potentially very extensive pleiotropic fitness target, one which can extend to the level of the entire social genome. Given that the majority of mutations are deleterious [15], the effects of mutations that act on the broader social genome are likely to be harmful to the fitness of populations and not merely individual carriers [16]. Thus the SEAM identifies a mechanism whereby individual carriers of certain mutations can influence the fitness characteristics of the social phenotype and genotype within which they are embedded. Although the SEAM was developed initially to explore the implications of deleterious variants accumulating under conditions of relaxed purifying selection in industrialized human populations on patterns of recent culture-gene co-evolution (see: Dutton et al., for a test of a prediction of this model [17]), a possible instance of this phenomenon was highlighted in the mouse utopia experiments of John Calhoun, wherein it was noted that the decline phase of Calhoun′s mouse populations co-occurred with an increasing prevalence of what he termed ″autistic-like″ behavioral phenotypes (″the beautiful ones″) [18]. Woodley of Menie et al. speculated that these ″beautiful ones″ might have established deleterious patterns of social epistasis, which had pleiotropic and disruptive effects at all levels of the mouse colonies′ social organization – engendering the colonies′ collapse [16].

Some studies have provided potential support for the SEAM, once again through analysis of mouse populations [3-5]. Crews et al. (2004) found that the distribution of social genotypes in the postnatal developmental environment of mice with a mutation affecting estrogen reception ″influenced[d] the development of sociosexual behaviors″ in these mice (p. 935). Similarly, Crews et al. (2009) found that the composition of mouse litters, in terms of sex ratios and relative frequencies of social genotypes, affected the ″aggressive behaviors″ of mice in adulthood (i.e. the genetic and sexual structure of the litters in which mice were raised had long-term developmental effects). Finally, Kalbassi et al. (2017) demonstrated the existence of disruptive IGE-patterns consistent with predictions of the SEAM in laboratory mice [3,19], showing that the absence of *Nlgn3,* a gene encoding the synaptic adhesion protein Neuroligin-3 and associated with Autism Spectrum Disorders [20-25], in male and female mice is sufficient to modify the behavioral phenotypes of their wild-type littermates in such a way that reduces certain of their fitness characteristics. Among the disruptive IGEs related to the presence of *Nlgn3* knockout mice were reduced testosterone levels and hierarchy-avoidant behaviors in mixed-genotype housings (i.e. housings containing knockout and wild-type mice). Kalbassi et al. (2017) also found evidence that social epistasis may promote phenotypic divergence and convergence in distinct traits; for example, *Nlgn3* knockout mice raised with wild-type littermates (in mixed genotype housing [MGH]) exhibited greater anxiety than knockout mice raised only with other knockout mice (in single genotype housing [SGH]), whereas wild-type mice were not significantly different with respect to anxiety across the two forms of housing. But in the case of social dominance and social interest, both wild-type and knockout mice showed depressed levels of these traits in MGH compared to SGH, thereby reducing the variance in this phenotype. Such findings indicate that IGEs may have a role in maintaining and, when compromised, degrading the integrity of group (specifically assemblage)-level fitness components, which potentially depend on the presence of certain levels of phenotypic variance with respect to certain group fitness optima [19]. We therefore postulated that in our experimental paradigm [3], defects in social hierarchy would be correlated with defects in social and territorial behaviors, two essential determinants of fitness. Furthermore, we hypothesized that IGEs would affect not only mouse behavior but also cause differential expression of genes associated with social communication.

## Results

The molecular mechanisms through which disruptive IGEs act are presently unknown. However, experiments conducted in *Drosophila serrata* and *melanogaster* showed that IGEs could modify the secretion of pheromones [6,12,13]. It is hypothesized that if such an effect is highly conserved across animal taxa, and is present in *Mus musculus,* it could constitute one element of the molecular and genetic pathway through which disruptive IGEs cause the social phenotype of the assemblage to shift away from its group-fitness optimizing state. *Nlgn3^y/+^* mice time spent more time with urine from unrelated *Nlgn3^y/+^* mice from SGH compared to urine from *Nlgn3^y/-^* mice from SGH (Figs 1A and B), suggesting that *Nlgn3^y/+^* mice urine composition differs from that of *Nlgn3^y/-^* mice. Major urinary proteins (MUPs) are pheromone proteins produced in the liver and excreted in the urine [26]. There are no identified post-translational modifications of MUPs and increased hepatic mRNA levels lead to increased secretion of the corresponding proteins in the urine [26]. Independently of the social environment, *Nlgn3^y/-^* mice show increased levels of *Mup4, Mup6* and *Mup20/Darcin* mRNA compared to *Nlgn3^y/+^* mice from SGH (Fig 1C). Experiments conducted in *Drosophila serrata* and *melanogaster* showed that IGEs could modify the secretion of pheromones [6,12,13]. Therefore we investigated the effect of social environment on the expression of hepatic *Mups. Nlgn3^y/+^* mice from MGH, born and raised with *Nlgn3^y/-^* mice as littermates, had greater levels of *Mup4, Mup6* and *Mup20/Darcin* mRNA compared to *Nlgn3^y/+^* mice from SGH (Fig 1C). By contrast, *Nlgn3^y/-^* mice from SGH and MGH expressed similar levels of hepatic *Mups* (Fig 1C). These results demonstrate that social environment has a selective effect on *Nlgn3^y/+^* mice in modifying their levels of hepatic *Mups.* Although social housing did not modify expression of hepatic sexual markers [3], it increased the expression of corticotropin release hormone receptor 2 *(Crhr2)* mRNA, a receptor controlling *Mups′* hepatic mRNA levels (supplementary Fig 1A) and overexpressed in socially submissive animals [27]. Lack of the dopamine D2 receptor (D2R) to control social hierarchy in neurons leads to a decrease in MUPs hepatic mRNA levels [28-30], so we hypothesized that dopamine signaling may control the increase in hepatic MUPs levels in *Nlgn3^y/+^* from MGH. Intraperitoneal injection of raclopride, a selective D2R antagonist, decreased *Mup20/Darcin* mRNA levels in the liver of mice from MGH but not of mice from SGH (Fig 1D). But injection of raclopride in *Nlgn3^y/+^* and *Nlgn3^y/-^* mice from SGH and MGH decreased the distance travelled in the open field to similar magnitudes (supplementary Fig 1B). These results suggest that MGH leads to a selective increase of activation of the hypothalamic-pituitary-adrenal axis by D2Rs, likely resulting in an increased CRHR2-dependent secretion of hepatic MUPs.

**Fig 1:**
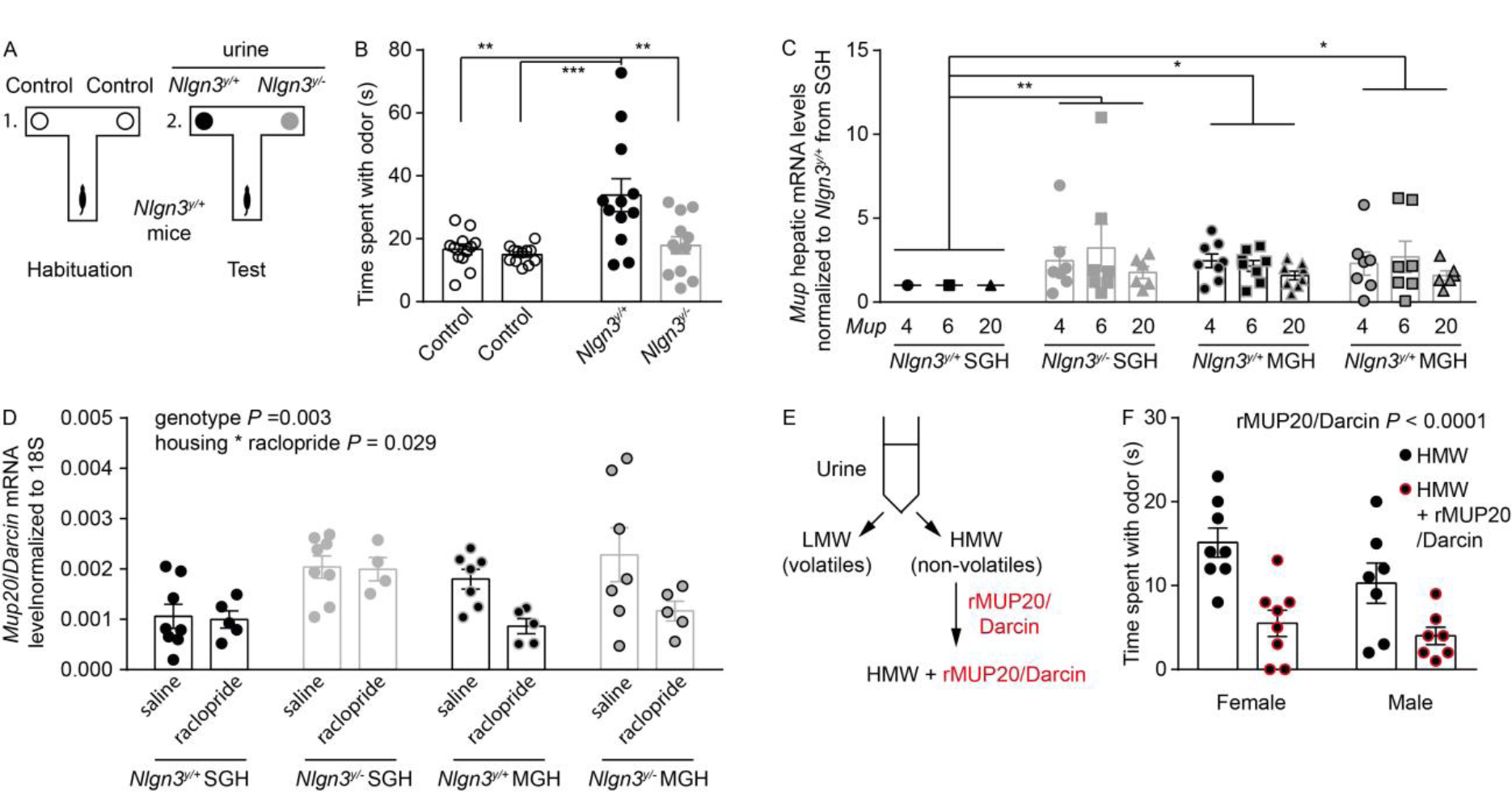
MUP20/Darcin produced by *Nlgn3^y/^′* mice is unattractive to *Ntgn3^Y/+^* mice. **(A)** Mice were placed in a T-maze and first habituated to control filter papers without odors and subsequently presented with two filter papers, one containing urine from *Nlgn3^y/+^* mice and the other containing urine from *Nlgn3^y/-^* mice. (B) *Nlgn3^y/+^* mice from SGH spent more time in contact with urine from *Nlgn3^y/+^* mice than urine from *Nlgn3^y/-^* mice (N = 12 *Nlgn3^y/+^* mice from SGH, Two-way ANOVA, genotype effect, *F*_(1,44)_ = 8.2, *P* = 0.0064, urine effect, *F*_(1,44)_ = 10.8, *P* = 0.002, interaction urine * genotype, *F*_(1,44)_ = 5.4, *P* = 0.025, Sidak′s post hoc test, ** *P* < 0.01 and *** *P* < 0.001). (C) Hepatic levels of *Mup4, 6* and *20* mRNA were lower in *Nlgn3^y/+^* mice from SGH compared to that of *Nlgn3^y/+^* mice from SGH and *Nlgn3^y/-^* mice from SGH and MGH (N = 8 *Nlgn3^y/+^* and *N* = 7 *Nlgn3^y/-^* mice from SGH and *N* = 8 *Nlgn3^y/+^* and *N* = 7 *Nlgn3^y/-^* mice from MGH, Two-way ANOVA, effect of housing, *F*_(1,78)_ = 6.3, *P* = 0.014, interaction housing * genotype, *F*_(1,78)_ = 4.15, *P =* 0.045, Sidak′s post hoc test, * *P*< 0.05 and ** *P*< 0.01). (D) Quantification of *MUP20/Darcin* mRNA levels in the liver of mice from SGH and MGH. *MUP20/Darcin* mRNA levels decreased 1 hour after injection of raclopride in mice from MGH but not in mice from SGH. (E) Urine was fractionated in components lower than 10kDa (low molecular weight – LMW) and higher than 10kDa (high molecular weight – HMW) and recombinant MUP20/Darcin (r MUP20/Darcin) was subsequently added to the HMW fraction. (F) *Nlgn3^y/-^* mice from SGH spent less time sniffing HMW fractions complemented with rMUP20/Darcin than HMW fractions without rMUP20/Darcin (N = 8 *Nlgn3^y/+^* mice from SGH, Two-way ANOVA, rMUP20/Darcin effect, *F*_(1,28)_ = 18.5, *P* = 0.0002). Values are represented as mean +/- S.E.M..

Among the pheromone proteins overexpressed by *Nlgn3^y/+^* mice from SGH, MUP20/Darcin, a pheromone protein expressed only by male mice [31], has the ability to trigger aggressive behavior [32]. We therefore speculated that increasing its levels in urine from *Nlgn3^y/+^* mice would be sufficient to reconstitute a *Nlgn3^y/-^-like* urine and decrease *Nlgn3^y/+^* mice interest. Male and female urines were purified to obtain low and high molecular weight (LMW and HMW) fractions containing volatile and non-volatile urinary (i.e. containing pheromone proteins) components respectively (Fig 1E) [33]. Addition of recombinant MUP20/Darcin (rMUP20/Darcin) to purified female and male MUPs led to a decrease in the time *Nlgn3^y/+^* mice from SGH spent with urinary pheromone proteins (Fig 1F), indicating that MUP20/Darcin is sufficient to decrease the interest of *Nlgn3^y/+^* mice from SGH in pheromone proteins and recapitulate the effect observed with urine from *Nlgn3^y/-^* mice.

Chronic exposure to MUPs during the development shapes mouse behavior [32]. We therefore hypothesized that *Nlgn3^y/+^* mice from MGH are chronically exposed to high levels of at least MUP4, 6 and 20 and, as a consequence, show different interest in pheromone proteins compared to *Nlgn3^y/+^* mice from SGH. Overall, *Nlgn3^y/+^* mice spent more time in interaction with social HMW fractions of male or female urine (Fig 2A and B, main effect of genotype *P* < 0.0001). This effect was due to *Nlgn3^y/+^* mice in SGH which overall spent more time with HMW fractions than *Nlgn3^y/+^* mice from MGH and *Nlgn3^y/-^* mice from SGH and MGH. Due to the overall difference in interest for pheromone proteins between genotypes and in order to avoid false positive results, we conducted the statistical analyses separately for *Nlgn3^y/+^* and *Nlgn3^y/-^* mice. *Nlgn3^y/+^* mice from SGH showed a greater time spent with HMW fractions of male and female urine compared to *Nlgn3^y/+^* mice from MGH (Fig 2A). In contrast to *Nlgn3^y/+^* mice from SGH, addition of rMUP20/Darcin to the HMW fraction of male and female urine did not change the time spent by *Nlgn3^y/+^* mice from MGH with urine fractions (Fig 2A). These results show that, when exposed to elevated concentration of rMUP20/Darcin, *Nlgn3^y/+^* from SGH show a decreased interest in HMW fractions of urine comparable to that of *Nlgn3^y/+^* mice from MGH. The addition of rMUP20/Darcin to HMW fractions of urine decreased the interest of *Nlgn3^y/-^* mice from MGH and SGH in these social odors (Fig 2B), suggesting that *Nlgn3^y/-^* mice can detect rMUP20/Darcin. The lack of interest in social cues observed in *Nlgn3^y/-^* mice from SGH could be caused by a chronic exposure to secreted MUP20/Darcin or an innate inability to detect specific social olfactory cues. To differentiate between the two hypotheses, we analyzed the behavior of *Nlgn3~^/^~* mice, as female mice do not express MUP20/Darcin. *Nlgn3^+/+^* mice showed a lower interest in female HMW urine fractions compared to *Nlgn3^y/-^* mice, similar to that of *Nlgn3^y/-^* and *Nlgn3~^/^~* mice (Fig 2C). MUP20/Darcin is an attractant for female mice [31]. *Nlgn3^+/+^* and *Nlgn3^-/^~* mice spent more time sniffing female HMW urinary fractions complement with rMUP20/Darcin than cotton only, demonstrating that they are both able to detect the male specific protein pheromone (Fig 2D). Nevertheless, *Nlgn3^-/-^* mice spent less time sniffing female HMW urinary fractions complemented with rMUP20/Darcin compared to *Nlgn3^+/+^* mice, suggesting that the lack of *Nlgn3* causes a decreased interest in rMUP20/Darcin and protein pheromones in general.

**Fig 2:**
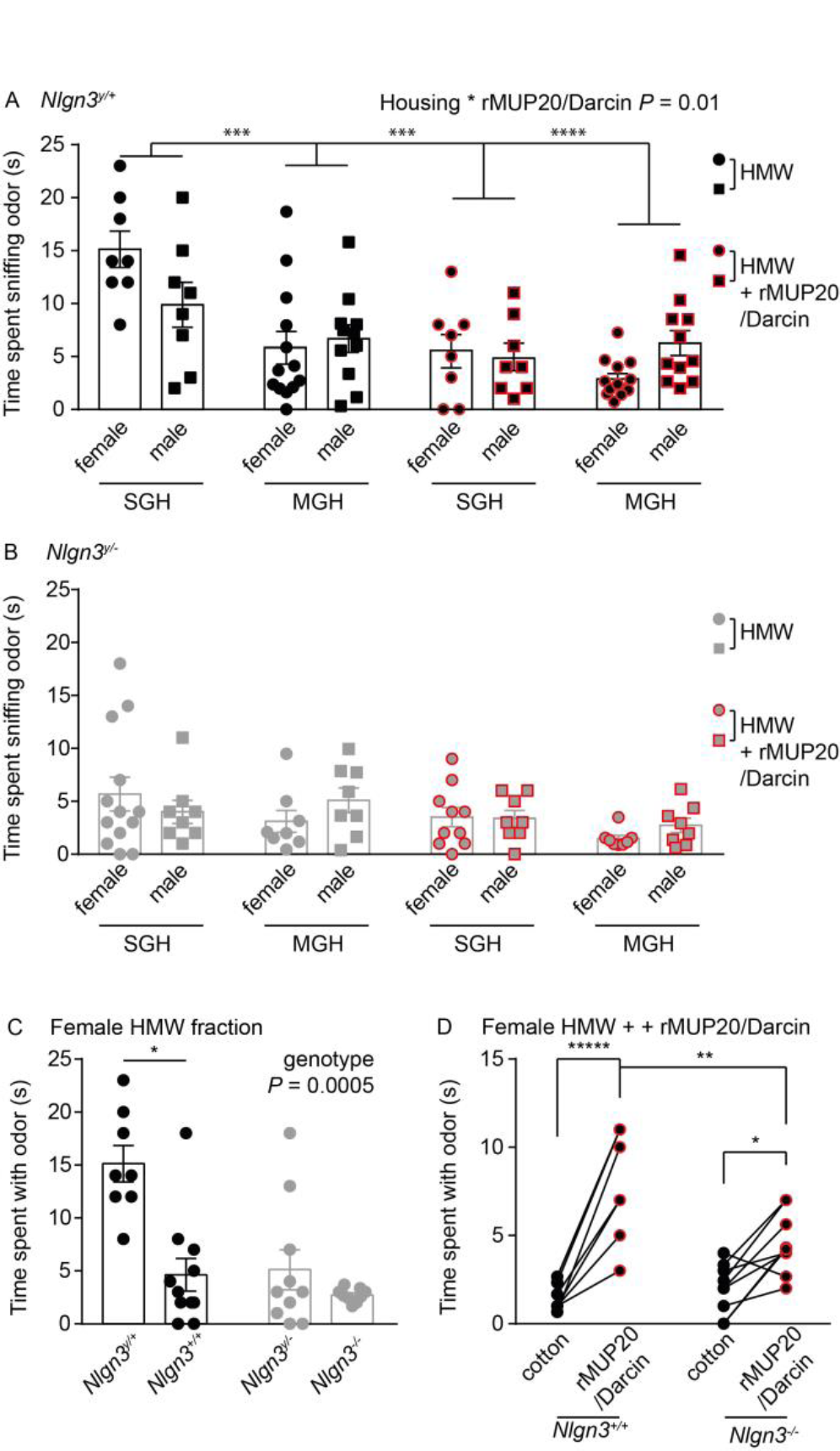
*Nlgn3^y/-^* mice modify *Nlgn3^y/+^* mice interest for protein pheromones. **(A)** *Nlgn3^y/+^* mice from SGH spent more time sniffing HMW fractions of male and female mice than *Nlgn3^y/+^* mice from MGH and than *Nlgn3^y/+^* mice from SGH and MGH sniffing HMW fractions of male and female supplemented with rMUP20/Darcin. As opposed to *Nlgn3^y/+^* mice from SGH, the addition of rMUP20/Darcin does not modify the interest of *Nlgn3^y/+^* mice from MGH for HMW fractions of urine (N = 8 *Nlgn3^y/+^* mice from SGH and *N* = 11 and 12 *Nlgn3^y/+^* mice from MGH, Two-way ANOVA, effect of rMUP20/Darcin, *F*_(1,70)_ = 21.0, *P* < 0.0001, effect of housing, *F*_(1,70)_ = 10.8, *P* = 0.0016, interaction rMUP20/Darcin * housing, *F*_(1,70)_ = 7.0, *P* = 0.001, Sidak′s post hoc test, *** *P* < 0.001 and **** *P* < 0.0001). Note that data related to *Nlgn3^y/+^* mice from SGH were replotted from Fig 1F. (B) *Nlgn3^y/-^* mice from SGH and MGH spent greater amount of time sniffing HMW fractions of male and female than HMW fractions of male and female supplemented with rMUP20/Darcin (N = 8-13 *Nlgn3^y/-^* mice from SGH and *N* = 8 *Nlgn3^y/-^* mice from MGH, Two-way ANOVA, effect of rMUP20/Darcin, *F*_(1,64)_ = 4.5, *P* = 0.0037). (C) *Nlgn3^y/+^* mice from SGH spent more time sniffing HMW fractions of female urine mice than *Nlgn3^+/+^* mice. Both *Nlgn3^y/-^* and *Nlgn3^/-^* mice spent similar amount of time sniffing HMW fractions of female mice urine compared to *Nlgn3^+/+^* mice but less compared to *Nlgn3^y/+^* mice (N = 8 *Nlgn3^y/+^* mice from SGH, *N* = 10 *Nlgn3^+/+^* mice, *N* =10 *Nlgn3^y/-^* mice from SGH and *N* = 10 *Nlgn3^-/^~* mice, Two-way ANOVA, effect of sex, *F*_(1,34)_ = 17.5, *P* = 0.0002, effect of genotype, *F*_(1,34)_ = 15.1, *P* = 0.0005, interaction sex * genotype, *F*_(1,34)_ = 7.0, *P =* 0.0126, Sidak′s post hoc test, * *P* < 0.05). (D) Both *Nlgn3^+/+^* and *Nlgn3~^/^~* mice spent more time sniffing HMW fractions of female mice urine complemented with rMUP20/Darcin. *Nlgn3^+/+^* mice spent more time sniffing HMW fractions of female mice urine complemented with rMUP20/Darcin than *Nlgn3~^/^~* mice (N = 7 *Nlgn3^+/+^* and *N* = 9 *Nlgn3~^/^~,* effect of rMUP20/Darcin, *F*_(1,14)_ = 56.0, *P* < 0.0001, interaction genotype * rMUP20/Darcin, *F*_(1,14)_ = 9.8, *P* = 0.0075, Sidak′s post hoc test, * *P* < 0.05, ** *P* < 0.01 and **** *P* < 0.0001). Values are represented as mean +/- S.E.M..

Discrimination between individuals of the same sex and typical courtship behavior require the ability to detect urinary protein pheromones [32,34]. When exposed to the two odors at the same time, *Nlgn3^y/+^* mice from SGH and from MGH spent more time sniffing the novel social odor compared to the familiar one (Figs 3A and B). Additionally, *Nlgn3^y/+^* mice from SGH and from MGH spent similar amounts of time in interaction with a female in estrus (Fig 3C). These results demonstrate that *Nlgn3^y/+^* mice from MGH have the ability to detect social cues but that their interest in them is decreased compared to that of *Nlgn3^y/+^* mice from SGH. *Nlgn3^y/-^* mice from SGH and MGH showed no interest in novel male social odors compared to familiar ones (Fig 3B) and spent less time with females in estrus compared to *Nlgn3^y/+^* mice from SGH and MGH (Fig 3C). Nevertheless, like *Nlgn3^y/+^* mice from MGH, *Nlgn3^y/-^* mice from MGH show a greater interest in novel female odors compared familiar male odors (Fig 3D).

**Fig 3:**
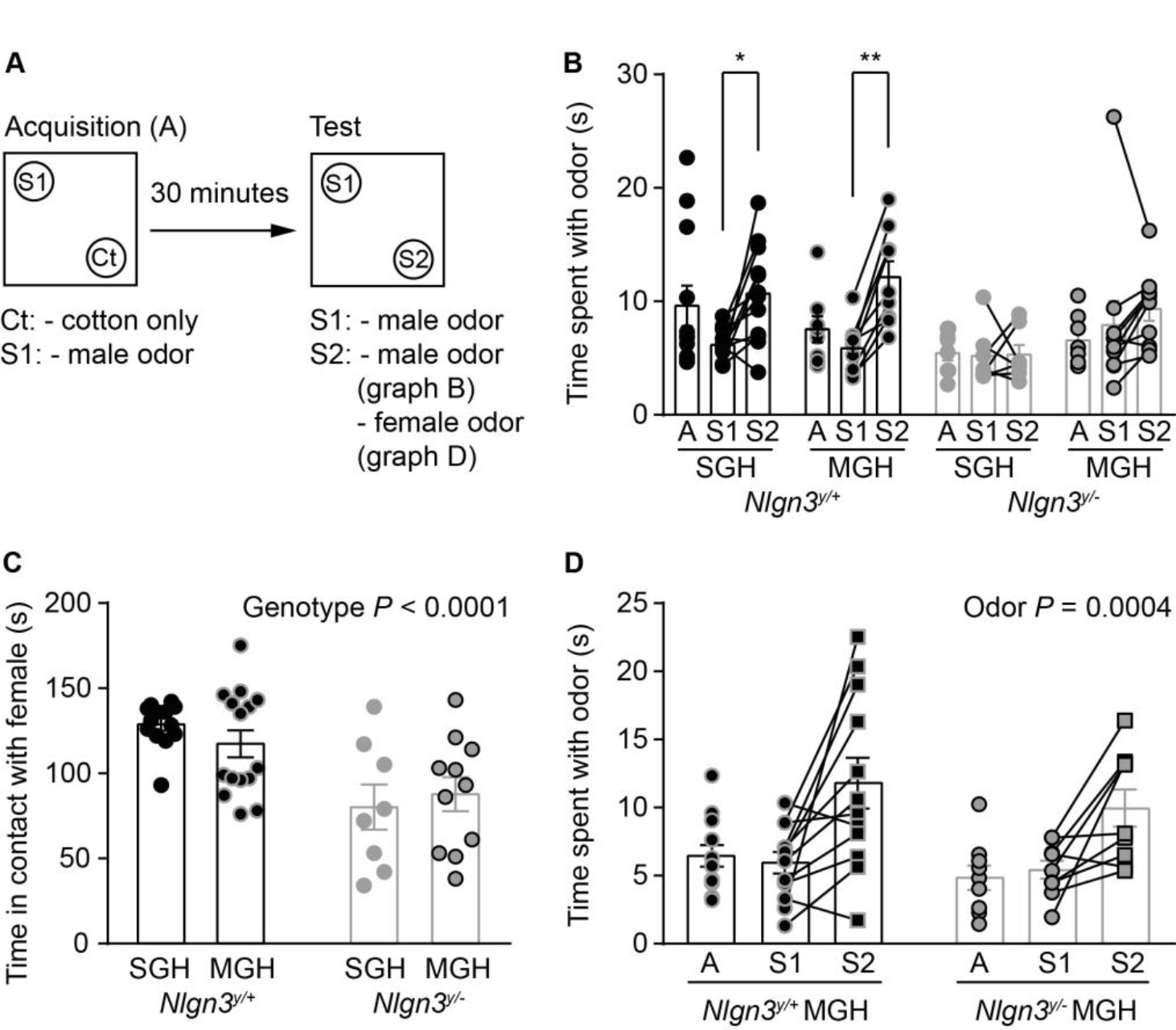
*Nlgn3^y/+^* mice show normal social discrimination and courtship behavior. **(A)** While in the dark, mice were first exposed to odors from a first set of males (S1) and to control. Thirty minutes after acquisition (A), mice were exposed to S1 and to odors from an unfamiliar set of males or females (S2). (B) *Nlgn3^y/+^* mice from SGH and MGH spent more time with S2 than with S1 during the test phase. *Nlgn3^y/-^* mice from SGH and MGH spent similar amounts of time with S2 and with S1 during the test phase. Note that *Nlgn3^y/+^* and *Nlgn3^y/-^* mice spent similar amount of time with S1 during the acquisition phase (N = 12 *Nign3^y/+^* and *N* = 7 *Nlgn3^y/-^* mice from SGH and *N* = 9 *Nign3^y/+^* and *N* = 10 *Nlgn3^y/-^* mice from MGH, Twoway ANOVA, effect of genotype, *F*_(1,68)_ = 4.0, *P* = 0.049, effect of housing, *F*_(1,68)_ = 5.0, *P* = 0.029, effect of odour, *F*_(1,68)_ = 12.1, *P* = 0.0008, interaction genotype * odour, *F*_(1,68)_ = 6.7, *P* = 0.0115, Sidak′s post hoc test, * *P* < 0.05 and ** *P* < 0.01). (C) *Nlgn3^y/+^* mice from SGH and MGH spent more time in contact with females in estrus than *Nlgn3^y/-^* from SGH and MGH (N = 13 *Nlgn3^y/+^* and *N* = 8 *Nlgn3^y/-^* mice from SGH and *N* = 15 *Nlgn3^y/+^* and *N* = 11 *Nlgn3^y/-^* mice from MGH, Two-way ANOVA, effect of genotype, *F*_(1,43)_ = 20.3, *P* < 0.0001). (D) *Nlgn3^y/+^* and *Nlgn3^y/-^* mice from MGH spent more time with S2 from females than with S1 from males. Note that *Nlgn3^y/+^* and *Nlgn3^y/-^* mice from MGH spent similar of time with S1 during the acquisition phase (N = 12 *Nlgn3^y/+^* and *N* = 9 *Nlgn3^y/-^* mice from MGH, Two-way ANOVA, effect of odour, *F*_(1,38)_ = 14.8, *P* = 0.0004). Values are represented as mean +/- S.E.M..

We demonstrated that IGE can impair the formation of social hierarchies in groups of mice from MGH by affecting their ability to compete for new territories but not their courtship behavior [3]. Territorial competition relies on both social and non-social behaviors and in particular on the ability to memorize new territories. The decreased social dominance observed in *Nlgn3^y/+^* and *Nlgn3^y/-^* mice from MGH could therefore be caused by a combination of social and non-social phenotypes. Mice from SGH habituate to new environments and, as a result, decrease their velocity between their first and second exposure to a new open-field environment (Fig 4A). Contrariwise, the distance travelled by mice from MGH did not decrease between day one and day two, showing a defect in memorizing a new environment. As we demonstrated previously, IGEs can be caused by a lack of *Nlgn3* in Pvalb-expressing cells [3]. Re-expression of *Nlgn3* in Pvalb-expressing interneurons of *Nlgn3^y/-^* mice was sufficient to restore a decreased distance travelled between day one and day two in both *Nlgn3^y/+^* and *Nlgn3^y/-^* mice, demonstrating that the deficit in memorizing new environments can be transferred through IGE. In our model system, we found that IGEs modify the behavior of group members and used Moore′s system [35] to describe our observations (Fig 4B). In this system, the phenotype of *Nlgn3^y/+^* mice in MGH (Z′_WT_) can be partitioned into three components: i) the genetic background (g_WT_), ii) the bidirectional IGE exerted by the hierarchical relationship between *Nlgn3^y/+^* and *Nlgn3^y/-^* mice Ψ_WT/KO_) [3] and iii) the unidirectional IGE exerted by MUP20/Darcin produced by *Nlgn3^y/-^* mice (Ψ_ko_) on the *Nlgn3^y/+^* mouse phenotype (ZWT, corresponding to the phenotype of *Nlgn3^y/+^* mice form SGH). The phenotype of *Nlgn3^y/-^* mice in MGH can be partitioned into two components: i) the genetic background (gKo) and the bidirectional IGE exerted by the hierarchical relationship between *Nlgn3^y/+^* and *Nlgn3^y/-^* mice (Ψ_WT/KO_) on the *Nlgn3^y/-^* mouse phenotype (Z_KO_, corresponding to the phenotype of *Nlgn3^y/-^* mice form SGH) [3].

**Fig 4:**
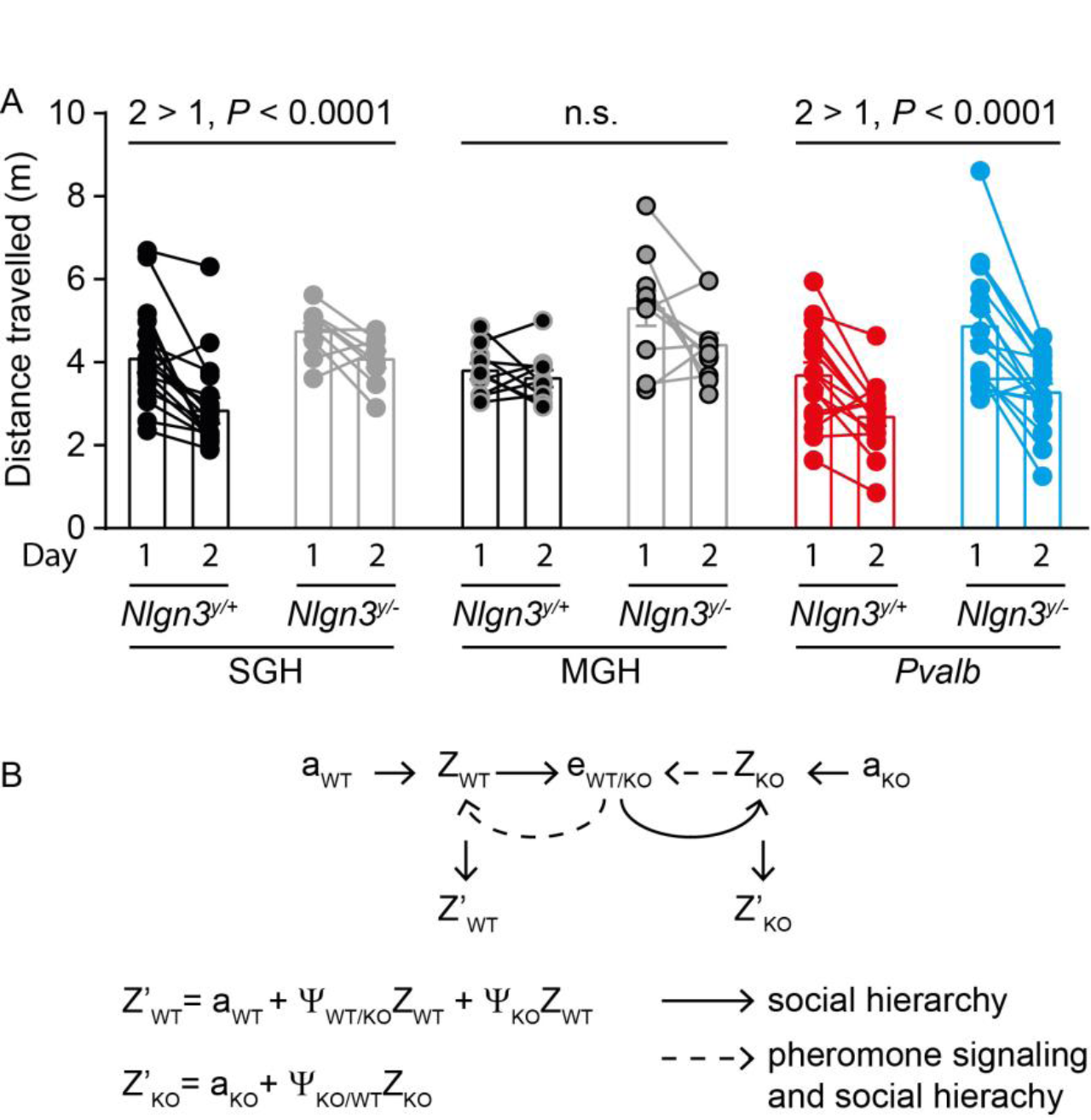
Theoretical framework for social epistasis in mouse models. **(A)** The activity of mice exposed to a new open-field environment for 20 minutes was recorded on day 1 and 24 hours after on day 2. The distance travelled by *Nlgn3^y/+^* and *Nlgn3^y/^-* mice from SGH and *Nlgn3^y/+^Pvalb^Cre^* and *Nlgn3^y/^-Pvalb^Cre^* mice on day 2 decreased compared to day 1. The distance travelled by *Nlgn3^y/+^* and *Nlgn3^y/-^* mice from MGH on day 2 was similar to that travelled on day 1 (N = 19 *Nlgn3^y/+^* and *N* = 14 *Nlgn3^y/-^* mice from SGH, *N* = 10 *Nlgn3^y/+^* and *N* = 10 *Nlgn3^y/-^* mice from MGH, *N* = 15 *Nlgn3^y/+^* and *N* = 11 *Nlgn3^y/-^* mice from *Pvalb,* Twoway ANOVA repeated measure, effect of genotype, *F*_(1,79)_ = 28.6, *P* < 0.0001, effect of housing, *F*_(2,79)_ = 3.4, *P* = 0.04, interaction day * housing, *F*_(2,79)_ = 3.1, *P =* 0.049, Sidak′s post hoc test, **** *p* < 0.0001). (B) Diagram illustrating the direct or genetic background (a) and indirect effects from the social environment (e) on the phenotype (Z) of *Nlgn3^y/+^* (WT) and *Nlgn3^y/-^* mice based on the equation of Moore et al.[35]. The phenotype of *Nlgn3^y/+^* mice (Z′_WT_) is a combination of genetic background with bidirectional (Ψ_WT/KO_Z_WT_) and unidirectional indirect genetic effects (Z_KO_) due caused by the social dominance of *Nlgn3^y/+^* mice and pheromone signaling respectively. The phenotype of *Nlgn3^y/-^* mice (Ψ_KO_Z_WT_) is a combination of genetic background with indirect genetic effects coming from the social environment (Ψ_WT/KO_Z_WT_), in particular their submissive behavior towards *Nlgn3^y/+^* mice. Values are represented as mean +/- S.E.M..

## Discussion

While substantial evidence of IGEs in a number of taxa has been documented, the molecular and genetic pathways by which they operate have remained unclear in most cases. This study demonstrates that in mice, exposure to the urinary pheromone protein MUP20/Darcin, which *Nlgn3^y/-^* mice excrete, alters the behavior of *Nlgn3^y/+^mice,* such that the latter exhibit reduced competitive behaviors. This phenotype is accompanied by a modification of the expression of hepatic *Mups* and *Crhr2* mRNA levels. These findings are consistent with the key prediction of the SEAM, which is that the negative fitness costs of deleterious mutations are not restricted to carrier organisms, but can extend to other organisms in carriers′ environments through modulation of gene expression. Our findings that pheromone protein expression is dependent on social context are consistent with results obtained in *Drosophila melanogaster* and *serrata* [6,12,13] and show that some basic mechanisms underlying IGEs might be conserved across animal taxa. Furthermore, we found that the increase in hepatic expression levels of *Mups* mRNA was due to a tonic activation of D2 receptor signaling, consistent with the role of D2R in controlling social hierarchy and protein pheromone secretion [28-30]. Therefore, central dopamine and hepatic MUPs systems represent both targets and vectors by which *Nlgn3^y/-^* mice exert IGEs on their wild-type littermates.

Mice carrying mutations associated with social disorders, which are inclined to higher levels of stress in the face of adversity than their wild-type counterparts, are more prone to react negatively to exposure to genetically unfamiliar mice, leading to phenotypic divergence with respect to anxiety between the two mouse types (this unidirectional effect is illustrated in Fig 4B). Both types of mice become socially avoidant in MGH because they are detecting conspecifics with which they are at genetic variance, which interferes with normal cues to hierarchy formation and social interaction, these being examples of adaptive processes that IGEs can disrupt when targeted by deleterious mutations (this bidirectional effect is illustrated in Fig 4B). Social aversion in both types of mice is taken on as a homeorhetic alternative stable state under conditions of such problematic genetic variation (and in the absence of corrective negative selection), further highlighting the role of certain kinds of mutations in affecting transitions between distinct alternative stable states to achieve metastability given variable population genetic composition; thus the MGH mice converge around a low-fitness behavioral phenotype characterized by social aversion (MGH mice likely have lower fitness, due to reduced competitive behaviors across the board, relative to SGH mice). These shifts in the behavior of wild-type and knockout mice have downstream negative effects on coordinative group-level processes that boost group fitness. The best evidence for such consequences is seen in the *de novo* memory loss phenotypes found in MGH in both wild-type and knockout mice. It is likely that when mice are unable to collectively map out, and therefore successfully navigate, their environment through urine pheromone secretion, they cannot acquire spatial memories of their surroundings, and must relearn the layout of their territory every time they move through it. As found in our experiments, these phenotypic alterations at the behavioral level are most likely underlain by modifications of genetic expression. In the current study, we identified alteration of genetic expression in the liver but we predict that they will also occur in cells in the central nervous system, and in *Pvalb-* expressing interneurons in particular [3].

Altogether, three IGE-related outcomes were observed: i) phenotypic divergence *(Nlgn3^y/-^* from MGH are more stressed than *Nlgn3^y/+^* and *Nlgn3^y/-^* from SGH), ii) phenotypic convergence *(Nlgn3^y/+^* mice adopt the disinterest in protein pheromones of *Nlgn3^y/-^* mice) and iii) *de novo* phenotypes in MGH (*Nlgn3^y/-^* and *Nlgn3^y/+^* mice lose their territorial memory and their socio-territorial competitive ability). Mechanisms leading to phenotypic convergence or to the emergence of *de novo* phenotypes are most likely causing the externalization of the fitness costs associated with the *Nlgn3^y/-^* mice phenotype (i.e. phenocopy) to their littermates. Such observations provide critical evidence that IGEs, when amplifying the effects of deleterious mutations, disrupt group-level processes that promote fitness (such as hierarchy formation). Since this disruption is seemingly triggered by the increased genetic variance in MGH compared to SGH – both wild-type and knockout mice are exposed to unfamiliar pheromones relative to SGH conditions – one possibility is that mice may be equipped with behavioral adaptations that dispose them to establish social bonds and groups with genetically similar conspecifics, and to avoid genetically dissimilar ones. Given the known role of Pvalb-expressing interneurons in mouse social behavior [3,36], it is possible that the behavioral systems undergirding these coalition-formation adaptations are associated with these neural structures. Consistent with this idea, the existence of a broader class of evolved behavioral regulatory mechanisms has been proposed, which promote a form of inter-organismal sociomonitoring that enable control of patterns of IGEs and group genetic architecture [37]. As previously noted, the fundaments of these mechanisms may be highly conserved across animal taxa, being present in *Drosophila,* a species in which pheromones play a role in regulating social interactions [2,6,12], but we anticipate that these mechanisms are far more sophisticated in species exhibiting highly complex sociality, and indeed may constitute the partial genetic underpinnings of human cultures [37].

## Methods

### Animals

All animal husbandry and experiments were performed in compliance with the UK Animals (Scientific Procedures) Act 1986, as amended, and in accordance with the Cardiff University animal care committee′s regulations. Mice containing a stop cassette flanked by loxP sites in the promotor region lacked *Nlgn3* expression *(Nlgn3^y/-^* #RBRC05451) [38] and mice expressing Cre recombinase under the *Pvalb* endogenous promoter in Pvalb-expressing cells *(Pvalb^Cre/Cre^* mice JAX:017320) [39] were backcrossed to a C57Bl/6 background for at least eight generations. Male and female mice were separated at weaning but housed only with their own littermates. For a summary of the breeding scheme, refer previous publication[3]. Sires were separated from pregnant dams and mice were weaned at postnatal day 30 to avoid the potential confounds associated with weaning on mice tested at postnatal day 21-28. Mice were kept on a 12-hour light/dark cycle with free access to food and water. All behavior was assessed during the light cycle. Experiments in adult mice were conducted when mice were 2-4 month old (Figs 1 to 3) and in young mice at postnatal day 21-28 (Fig 4). To minimize anxiety associated with human handling, all mice were well handled prior to testing [40]. On the testing day mice were habituated for 30 minutes to the testing room and handled with minimal restraint to reduce anxiety[40]. Note that all mice did not undergo testing in all tasks.

### Interaction with females

Prior to experiments, vaginal smears were stained with modified giemsa solution (fixative and blue/azure dye) to determine the stage of estrus cycle [41]. Test male mice were first habituated for 3 minutes to the arena. Subsequently, an unfamiliar female mouse in estrous was added to the same arena for 3 minutes. An experimenter blind to genotype manually scored interaction times. Interaction was recorded when mice were within 2 centimeters of each other.

### Interest in social odors

Urine was taken from adult male SGH *Nlgn3^y/-^* or *Nlgn3^y/+^* mice immediately after bladder voiding and frozen until use. 10µl of urine was pipetted onto blotting paper for experimental use (Fig 1A). A T-maze was used to assess time spent with the urine of *Nlgn3^y/-^* and *Nlgn3^y/+^* mice. SGH *Nlgn3^y/+^* mice were placed into the long arm of the T-maze and allowed to freely explore the T-Maze. Two trials were conducted, the first being a habitation period in which no urine on blotting paper was administered. The second being the trial, in which the urine on the blotting paper was placed in opposite arms. The sides in which the urine was placed were altered to avoid bias. The trial started when the mouse was put into the long arm of T-maze, and was allowed to freely explore. Each trial lasted 2 minutes each. EthoVision XT tracking software (Noldus) was used to record the activity of the mice within the T-Maze. The interest to the odour was expressed as time, in seconds, spent with the odour.

Urine from several *Nlgn3^y/+^, Nlgn3^+/+^* or *Nlgn3^y/-^* mice from SGH was collected and pooled to produce a mixture of samples from different home cages. Fractionation of urine was performed following previously published methods [33] (Fig 1E). Briefly frozen urine pooled for each genotype was kept on ice then fractionated using Amicon^®^Ultra-15 centrifugal filters (Millipore) to produce low molecular weight (LMW) urine faction and high molecular weight (HMW) urine fraction. The LMW fraction was collected in the flow through and the HMW fraction was removed from the filter by dilution to original total volume with artificial urea (NaCl 120 mM, KCl 40 mM, NaH4OH 20 mM, CaCl2 4 mM, MgCl2 2.5 mM, NaH2PO4 15 mM, NaHSO4 20 mM, Urea 333 mM at pH 7.4).

For social interest experiments (Fig 1F and 2A-D) mice were placed in an arena with a hole in one side for introducing stimuli on cotton swabs. Mice were acclimatized to the arena for two minutes and then to the presence of a clean cotton swab for two minutes prior to testing. Following a one minute break (where no cotton swab was presented), mice were exposed to a HMW fraction complemented or not with recombinant MUP20/Darcin [31,42] (gift from Prof Hurst, Liverpool University). Each mouse was exposed to each of the stimuli over a series of days (order of odor exposures were counterbalanced).

HMW urine fractions were administered at physiological concentration (10 μl per swab) with or without recombinant MUP20/Darcin (0.2ug/ml). Behavior was scored as time spent interacting (sniffing, biting etc.) on the portion of the cotton swab which contained the scent cue, interactions with the stick of the swab (leaning, climbing etc.) were not included.

For social discrimination experiments (Fig 3B and D), the social odors consist of cage scrapings originating from two cages of three C57Bl/6 male or female mice maintained for 6 days with the same home cage bedding to allow for a concentration of odorants. During the acquisition phase, a swab containing a social odor (S1) and a swab without odor were presented in two identical cups placed in opposite corners of the open field for 10 minutes. Mice were placed back in their home cage for 30 minutes and subsequently exposed to a swab containing S1 and a swab containing the odor of unfamiliar male or female mice (S2) in the same open field. Mice were able to be in direct contact with the odors. The time spent in proximity to the social odor (<10 centimeters from the swab) was quantified using EthoVision XT^®^ tracking software (Noldus, Wageningen Netherlands).

### Spontaneous activity

Spontaneous activity of mice was recorded in a 40 centimeter x 20 centimeter open field arena for 20 minutes in the dark using an infrared video camera over two consecutive days. EthoVision XT^®^ tracking software (Noldus, Wageningen Netherlands) was used to measure the distance traveled in the OF (average meters traveled).

### RNA isolation and quantitative Real-time PCR

Total RNA from liver was isolated with TRIzol^®^ reagent (Thermo Fisher Scientific, Carlsbad, California) and purified using the RNeasy kit (Qiagen GmbH, Hilden, Germany). The cDNA was synthesized using Superscript III (Thermo Fisher Scientific, Carlsbad, California). Quantitative real-time PCR analysis was performed using Fast SYBR green Master Mix^®^ (Thermo Fisher Scientific, Vilnius, Lithuania) on a real-Time PCR System (Thermo Fisher Scientific, Carlsbad, California). Relative expression levels were determined by normalization to 18S rRNA expression using the comparative ∆∆C_T_ method. Primers used: MUP4 forward: 5′- ATGAAGCTGCTGCTGTGT -3′; MUP4 reverse: 5′- TCATTCTCGGGCCTTGAG -3′, MUP6 forward: 5′- ATGAAGCTGCTGCTGCTGT -3′; MUP6 reverse 5′- TCATTCTCGGGCCTGGAG -3′; MUP20 forward 5′- CTGCTGCTGTGTTTGGGACT -3′; MUP20 reverse 5′- TCTTTTGTCAGTGGCCAGCA -3′; CRHR forward 5′- GCATCACCACCATCTTCAAC -3′; CRHR2 reverse 5′- GAATGCACCATCCAATGAAG -3′; 18S forward: 5′-GTCTGTGATGCCCTTAGATG-3′; 18S reverse: 5 ′ – AG CTTATGACCCGCACTT AC-3 ′.

### Statistical analysis

We designed our tests groups to have mice from the different groups tested at the same time. We used GraphPad Prism^®^ to systematically test for normality using the D′Agostino-Pearson test, and for outliers using the ROUT method with *Q* = 1 to ensure no outliers would modify the outcome and power of the statistical tests. No animals were removed from the analyses. For each experiment, at least three independent litters were analyzed. Multiple comparisons were performed using Two-way non-repeated and repeated measure analysis of variance (ANOVA) as all our datasets followed a normal distribution, and, when appropriate, followed by post-hoc Sidak′s test.

## Acknowledgments

We thank J. Hurst for providing recombinant MUP20/Darcin. Funding sources: This work was supported by the Life Science Research Network Wales, an initiative funded through the Welsh Government′s Ser Cymru program and by funds to S.J.B and E.C. from the Wellcome Trust, UK. The authors declare no conflict of interest. S.O.B., E.C. and S.K. performed research, analyzed data and participated in writing the manuscript. M.A.S. and M.A.W. wrote the manuscript. S.J.B. designed and performed research, analyzed data and wrote the manuscript.

Authors report no conflict of interest

---

**Supplementary Fig for Fig 1:**
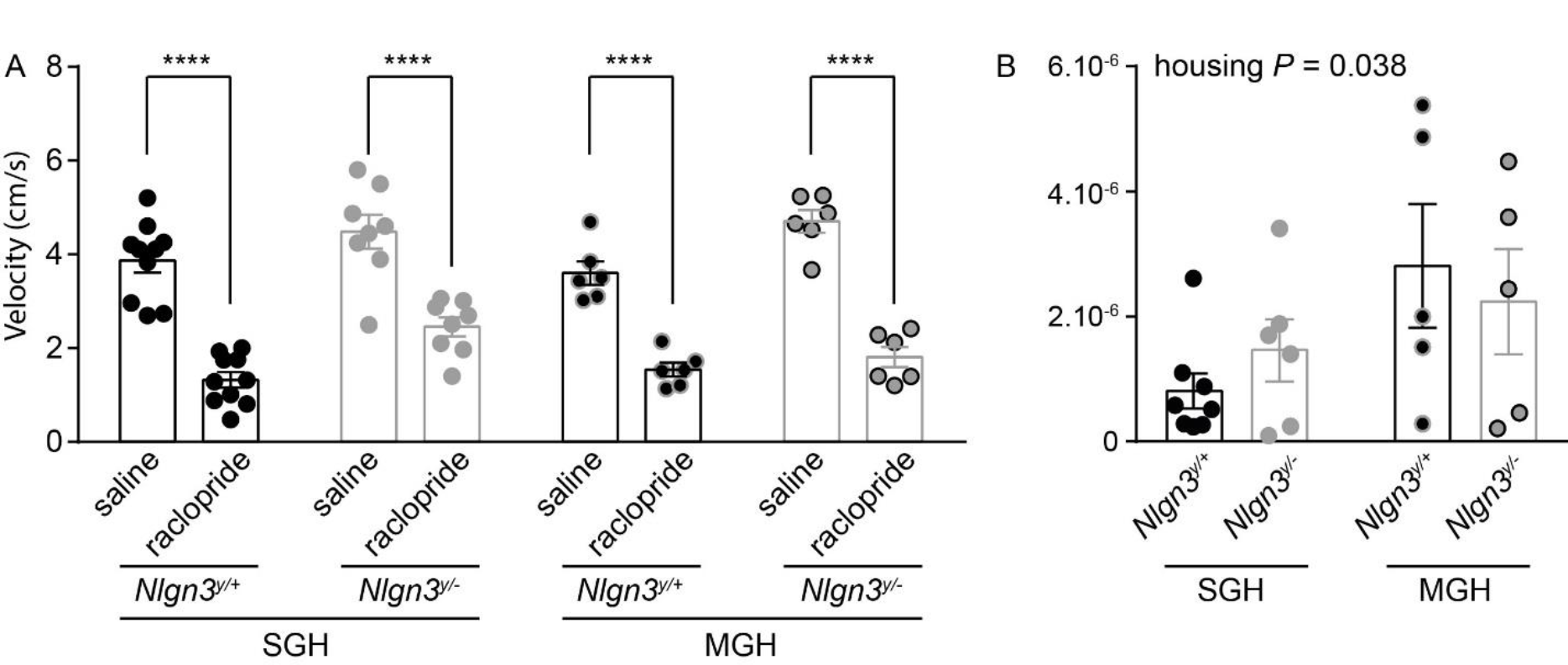
**(A)** Independently of the housing condition, injection of raclopride leads to a decrease of velocity in *Nlgn3^y/+^* and *Nlgn3^+/−^* mice (N = 10 *Nlgn3^y/+^* and *N* = 8 *Nlgn3^y/-^* mice from SGH and *N* = 6 *Nlgn3^y//^+* and *N* = 6 *Nlgn3^y/-^* mice from MGH, Two-way ANOVA, effect of raclopride, *F*_(1,52)_ = 176.6, *P* < 0.0001, effect of genotype *F*_(3,52)_ = 6.948, P = 0.0005, Sidak′s multiple comparison, **** *p* < 0.0001). (B) *Nlgn3^y/+^* and *Nlgn3^y/-^* mice from MGH have increased levels of hepatic CRHR2 mRNA compared to *Nlgn3^y/+^* and *Nlgn3^y/-^* mice from SGH (N = 8 *Nlgn3^y/+^* and *N* = 6 *Nlgn3^y/-^* mice from SGH and *N* = 5 *Nlgn3^y/+^* and *N* = 5 *Nlgn3^y/-^* mice from MGH, Two-way ANOVA, effect of housing, *F*_(1,20)_ = 4.9, *P* = 0.038).

